# Host genetic and environmental factors shape the human gut resistome

**DOI:** 10.1101/2020.05.18.092973

**Authors:** C.I. Le Roy, R.C. E. Bowyer, V.R. Carr, R. Costeira, J.E. Castillo-Fernandez, T.C. Martin, T.D. Spector, C.J. Steves, D. Moyes, S.K. Forslund, J.T. Bell

**Author notes:** Equal contribution (S.K. Forslund, J.T. Bell). Correspondence to Sofia Forslund, and Jordana Bell.

## Abstract

**Background:** Understanding and controlling the spread of antimicrobial resistance is one of the greatest challenges of modern medicine. To this end many efforts focus on characterising the human resistome or the set of antibiotic resistance determinants within the microbiome of an individual. Aside from antibiotic use, other host environmental and genetic factors that may shape the resistome remain relatively underexplored.

**Methods:** Using gut metagenome data from 250 TwinsUK female twins, we quantified known antibiotic resistance genes to estimate gut microbiome antibiotic resistance potential for 41 types of antibiotics and resistance mechanisms. Using heritability modelling, we assessed the influence of host genetic and environmental factors on the gut resistome. We then explored links between gut resistome, host health and specific environmental exposures using linear mixed effect models adjusted for age, BMI, alpha diversity and family structure.

**Results:** We considered gut microbiome antibiotic resistance to 21 classes of antibiotics, for which resistance genes were detected in over 90% of our population sample. Using twin modelling, we estimated that on average about 25% of resistome variability could be attributed to host genetic influences. Greatest heritability estimates were observed for resistance potential to acriflavine (70%), dalfopristin (51%), clindamycin (48%), aminocoumarin (48%) and the total score summing across all antibiotic resistance genes (38%). As expected, the majority of resistome variability was attributed to host environmental factors specific to an individual. We compared antibiotic resistance profiles to multiple environmental exposures, lifestyle and health factors. The strongest associations were observed with alcohol and vegetable consumption, followed by high cholesterol medication and antibiotic usage. Overall, inter-individual variation in host environment showed modest associations with antibiotic resistance profiles, and host health status had relatively minor signals.

**Conclusion:** Our results identify host genetic and environmental influences on the human gut resistome. The findings improve our knowledge of human factors that influence the spread of antibiotic resistance genes and may contribute towards helping to attenuate it.

## Background

Currently, antibiotics are the most effective treatment for infectious diseases in humans and in animals. However, their intensive use in health care and food production has led to a dramatic increase in antibiotic resistant pathogens [1]. Antibiotic resistance is acquired by bacteria through mutation and gene transfer. The human gut is home to trillions of bacteria and can act as a reservoir for antibiotic resistance genes (ARG), where exchange of ARG may take place between bacteria [2, 3]. ARG can be transferred vertically throughout bacterial division and horizontally between bacteria *via* transformation (integration of DNA fragments from the environment), transduction (through a bacteriophage) and conjugation (interaction between two bacteria) [4]. Individuals are constantly exposed to new bacteria that might reach the gastrointestinal track and although the ability of these bacteria to colonise the large intestine is debated [5], their passage through the gut ecosystem may be sufficient to horizontally transfer ARGs to the microbial community. Thus, the host microbiome may have the potential to acquire antibiotic resistance without direct antibiotic exposure. Resistant pathogenic bacteria are a serious health problem, and resistant non-pathogenic bacteria are also of concern due to their potential to transfer ARGs to pathogens. Indeed, the continuous rise of antibiotic resistant bacteria has led to a significant increase in mortality, especially in nosocomial infections [6].

Advances in technology have allowed for the collective sequencing of whole gut microbiota genomes, or metagenomes [7]. It is therefore possible to identify and potentially quantify ARG carried by bacteria in the gut community through the analysis of gut metagenome data. Several studies have explored the ARG profile of the human gut microbiome [8, 9], or the gut resistome, using different approaches including total number of ARGs in the gut or metrics such as the antibiotic resistance potential (ARP) [10]. ARP estimates the number of ARG copies in a sample, weighted by the relative abundance of taxa carrying the ARG. Although ARP metrics do not measure functional antibiotic resistance, they have been used to explore factors that may shape the gut resistome. For instance, significant ARP differences were observed across countries mirroring differences in country-specific antibiotic consumption [11], where higher antibiotic use in human, and also farm animals, was related to greater ARP levels. Medicinal antibiotic use plays an important role in shaping the gut resistome, where antibiotic use during hospitalisation has been associated with increased relative abundance of ARGs in the gut [12]. The potential for transition of ARGs during food production, or the ‘farm-to-fork’ hypothesis, has been extensively discussed in the literature [13]. Although evidence remains sparse [14, 15], direct exposure to livestock has been linked to an increase in the number of ARG within the human gut [16]. Furthermore, other environmental or lifestyle factors have also been linked to gut resistome variation [17]. For example, significant gut resistome associations with travel [18] and pet ownership [19] suggest that a multitude of factors could be at play.

Despite this, the factors shaping the human gut ARG reservoir are still not well understood. Exploring country-specific environmental variation allows insight into environmental parameters involved in this process [8, 10, 20]. In addition, previous work has demonstrated that the gut microbiome could also be influenced by host genetics [21, 22], with even stronger influence observed when considering gut microbiota fonctionality [23]. Therefore, it is plausible that host genetic impacts may also affect the abundance of bacteria that carry ARGs, as well as the potential to transfer ARGs in the gut community.

In this study, we hypothesised that both host genetic and environmental factors influence the human gut ARG reservoir. By profiling the ARP in a sample of 250 healthy female volunteers from the TwinsUK cohort, we evaluated the role of host genetic and environmental impacts on the resistome using a twin study design. We then explored resistome associations with specific environmental factors and health status in shaping the human gut ARG reservoir.

## Methods

### Samples

We used published gut metagenomic profiles of 250 female twins from the TwinsUK cohort of mean age 61 (range 36-80 years of age). The sample contained 35 monozygotic (MZ) and 92 dizygotic (DZ) twin pairs with an average body mass index (BMI) of 25.8 ± 4.61. Sample collection and sequencing methods have previously been described [23], with on average 74 million non-human high-quality Illumina HiSeq paired-end reads of a read length 100 bp (insert size 350 bp) per sample. Sequence data quality control, gene catalogue build, gene abundance estimation, and taxonomic assignment have previously been described in this dataset [23]. Briefly, the published gene catalogue consisted of 11,446,577 non-redundant genes, at which relative gene abundances were estimated [23] using previously described methods [24, 25]. Taxonomic annotation has previously been described in this sample [23], and utilised taxonomic assignments from the IGC gene catalogue [26] and application of the same pipeline [25, 25] for taxonomic assignment of the additional genes reported in this sample [23]. The relative abundance of a taxon is calculated from the relative abundance of its genes, considering only signals with at least 10 genes from a taxon.

### Antibiotic Resistance Potential

Gut resistomes were profiled using the antibiotic resistance potential (ARP) approach [10]. The ARP is defined as the average microbial genome fraction encoding ARGs for a particular antibiotic or class of antibiotics, across all bacteria in the gut microbiome sample, based on known taxonomy of the ARGs (here considered at the genus level, with each genus represented by its average ARG carriage within the ProGenomes database) [27]. The approach uses the above described gene catalogue, published relative gene abundances and catalogue amino acid sequences to assess ARG abundance in the sample and subsequently takes into account their taxonomic composition to generate the ARP. For ARP estimation amino acid sequences were translated from the gene sequences, selecting the frame resulting in the longest uninterrupted protein, and where for the majority of sequences (80%) only one specific frame was full length and was selected. The gene catalogue in this dataset was the annotated for ARGs using CARD (version 2.0.1) [28] and ResFams [29], assigning ResFams hits only to sequences without a CARD hit and integrating both types of annotation via the Antibiotic Resistance Ontology (ARO). This resulted in a gene catalogue annotated with ARG family membership and thus total gene abundances per ARG family. Together with projections on expected ARG abundance from taxonomic composition of each sample, ARPs were then computed. The ARP is a measure of antibiotic resistance gene abundance relative to the amount of sample material stemming from taxa known to carry such resistance genes. The measure aims to decouple ARGs increases following from taxonomic composition change only, compared to changes resulting from selection within taxonomic groups for higher ARG carriage. Thus, findings of altered raw ARG abundance versus altered ARP abundance represent different scenarios each leading to altered resistance capacity in microbial ecosystems – ARG shift in the absence of ARP shift reflects changes driven by larger-scale taxonomic composition shift with accompanying changes in ARG abundance, whereas ARP shift may indicate a shift within taxa to more resistant varieties, including by direct propagation of resistance genes, copy number alterations, mobile element transmission, strain replacement or other scenarios. ARP were estimated for 339 profiles that clustered and represented resistance to 39 specific types of antibiotics or classes of antibiotics, many of which were highly correlated. Altogether, estimates were obtained for 41 different variables, spanning 39 types of antibiotics or classes of antibiotics, one antibiotic resistance mechanism represented as a proxy class (efflux pumps), and the overall total of resistance genes within an individual. Pair-wise correlations were estimated across the 41 variables, with multiple highly correlated profiles (**Supplementary Figure 1**). Therefore, a single ARP was chosen to represent each cluster of highly correlated of ARPs (pair-wise Spearman rho > 0.9), selecting the most prevalent profile as the representative per correlated cluster (**Supplementary Figure 1**). ARP profiles were then corrected for potential covariates in a linear mixed effects regression to generate the ARP residuals that were included in the majority of downstream analyses. Covariates included BMI, age, and α-diversity as fixed effects, and family and zygosity as random effects.

### Twin modelling: ARP heritability and environment effects

Twin-based heritability of ARP variables was calculated by fitting the ACE model to ARP residuals using the ‘OpenMx’ package in R version 3.6.1. The model assesses the relative contribution of additive genetic effects (A), common environment (C), and environment unique to an individual (E), towards the variance of a phenotype of interest, here a specific ARP residual profile (http://openmx.ssri.psu.edu/docs/OpenMx/2.3.1/GeneticEpi_Path.html). The significance of the A component was based on the difference between the fit of the ACE and the CE models to evaluate if inclusion of A fit the data better than use of C and E alone.

### Association study

To follow up twin-based results of host environmental influences on ARPs, we carried out association analyses comparing inter-individual variability in each ARP profile to a series of host environmental variables. Host environmental variables included factors related to health and host environment, such as lifestyle and diet factors, and medication use. We identified 24 health markers that included 21 conditions that were reported in at least 10 of the 250 twins, as well as further variables such as number of days spent in a hospital, DEXA measures of visceral fat mass, and estimated frailty [30] and neuroticism scores [31] (**Supplementary table 1)**. Next, a total of 32 environmental factors were selected and divided in three categories: distal environment (5), diet (14) and medication use (12). Information related to diet, lifestyle, medication use, and health status were collected through questionnaires sent to the volunteers and time matched with the date of sample collection. Dietary intakes were estimated via food frequency questionnaire (FFQ) data, collected following Epic-Norfolk guidelines [32], and used to construct the Healthy Eating Index (HEI) 2010 [33], previously validated within this cohort as a means of capturing dietary variance [34]. The index of multiple deprivation (IMD), a composite measure of area-level deprivation, was downloaded from government websites and used to derive within-population quintiles as described previously [35]. Environmental data were not always available for the 250 twins and details of the sample size for each variable can be found in **Supplementary table 1**.

To evaluate the association between each individual ARP and the environmental variable of interest we used a linear mixed effects regression model (lme4 package in R version 3.6.1). Unadjusted ARPs were fit as the response variable, the environmental or health variable was the predictor, and models were adjusted for BMI, age, alpha diversity and family structure as previously described. Significance of the results were evaluated by comparing the full model (including the variable of interest) to a null model (excluding the variable) using a likelihood ratio test. Results were adjusted for multiple testing using the false discovery rate (FDR 5%).

## Results

### Profiling the gut resistome

We explored gut metagenomic profiles of 250 healthy older (mean age 65) Caucasian female twins from the TwinsUK cohort, including 35 MZ and 92 dizygotic DZ twin pairs. The gut resistome in each individual was characterised using the antibiotic resistance potential (ARP), a previously developed measure of ARG abundance relative to abundance of their likely carrier taxa [10]. ARP profiles were estimated for 41 variables, which included antibiotics, antibiotic classes, and antibiotic resistance mechanisms. Some of the ARP variables were highly correlated and therefore replaced by the most prevalent profile as a representative of each cluster (**Supplementary Figure 1**). Altogether, 23 ARP profiles were less correlated and therefore considered as independent variables (Spearman rho < 0.9). The variables assess potential of AR to specific antibiotics and classes of antibiotics, including ARP for the total gut resistome estimated as the overall sum of ARPs within an individual, or total ARP (ALL). Of the 23 ARPs, 21 were detected in over 90% of our sample and were explored in subsequent analyses (**Figure 1**). Therefore, in most of our UK population sample the gut communities could be considered as carriers of a large proportion of well characterised ARGs (**Figure 1**). Tetracycline and clindamycin - two broad spectrum antibiotics widely used in humans - were the ARPs detected at the highest level in our sample (**Figure 1**). In contrast, ARPs to amythiamicin A and fusidic acid were detected in less than 20% of the population sample and were excluded in downstream analyses in this study.

**Figure 1:**
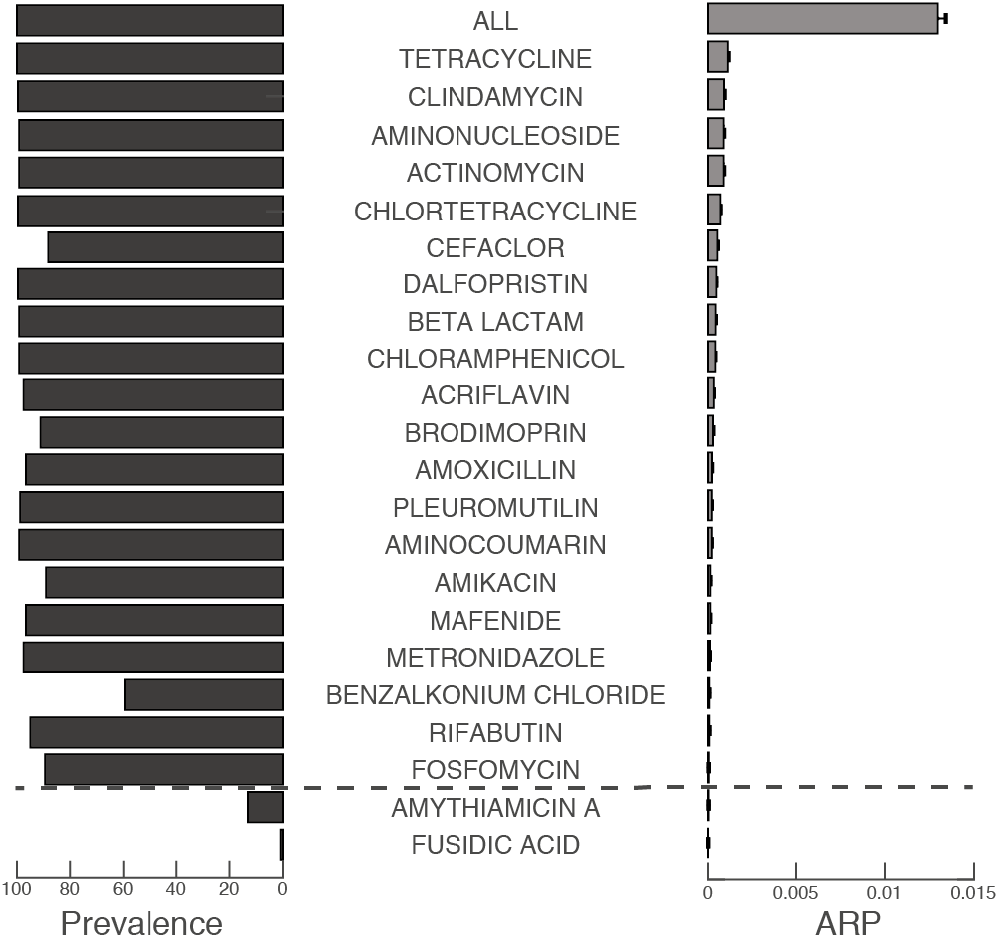
Antibiotic resistance potential level and prevalence among the TwinsUK cohort. Prevalence among the population is pictured on the left and mean ARP levels on the right. ARPs below the dotted line are removed from subsequent analyses.

### Host genetic influences on the gut resistome

Since our samples constitute only twin pairs, we carried out twin-based heritability analyses of the ARP profiles. Using the ACE model, we estimated the proportion of variation that is attributed to host genetic or environmental factors for each of the 21 ARP variables.

We observed that ARP profiles are predominantly under the influence of host environmental factors (**Figure 2A, Supplementary table 2**). However, two ARP profiles had strong evidence for heritability (A > 50%), namely acriflavin (A = 70%, 95% CI = [36-85]%) and dalfopristin (A = 51%, 95% CI = [6-72]%). Altogether, five ARPs displayed a nominally significant fit of the heritability term in the twin model, and these were acriflavin, dalfopristin, aminocoumarin (A = 48%, 95% CI = [1-69]%) and clindamycin (A = 48%, 95% CI = [4-71]%), as well as total ARP (ALL, A = 38%, 95% CI = [1-65]%). In total 12 ARPs (57% of profiles) had at least modest heritability estimates over 20% (A > 20%). The average ARP heritability across the 21 variables was estimated to be over 25% (A = 28.4% ± 21.4, **Figure 2A**). The four ARPs displaying greatest heritability estimates (acriflavine, dalfopristin, aminocoumarin, clindamycin) were highly prevalent in our sample (>95% prevalence, **Figure 1**) and in an independent gut metagenomic dataset from healthy Western Europeans (>=50% prevalence of cluster components in Carr et al. 2020 [20], **Supplementary table 3**). We also verified that highly correlated ARPs (Spearman rho > 0.9) displayed similar levels of heritability estimates (**Figure 2B**). For instance, acriflavine that was the most heritable ARP (A = 70%) and was highly correlated with four other ARP measures (ciprofloxacin, moxifloxacin, nalidixic acid and norfloxacin) that all displayed nominally significant heritability estimates above 50%.

**Figure 2:**
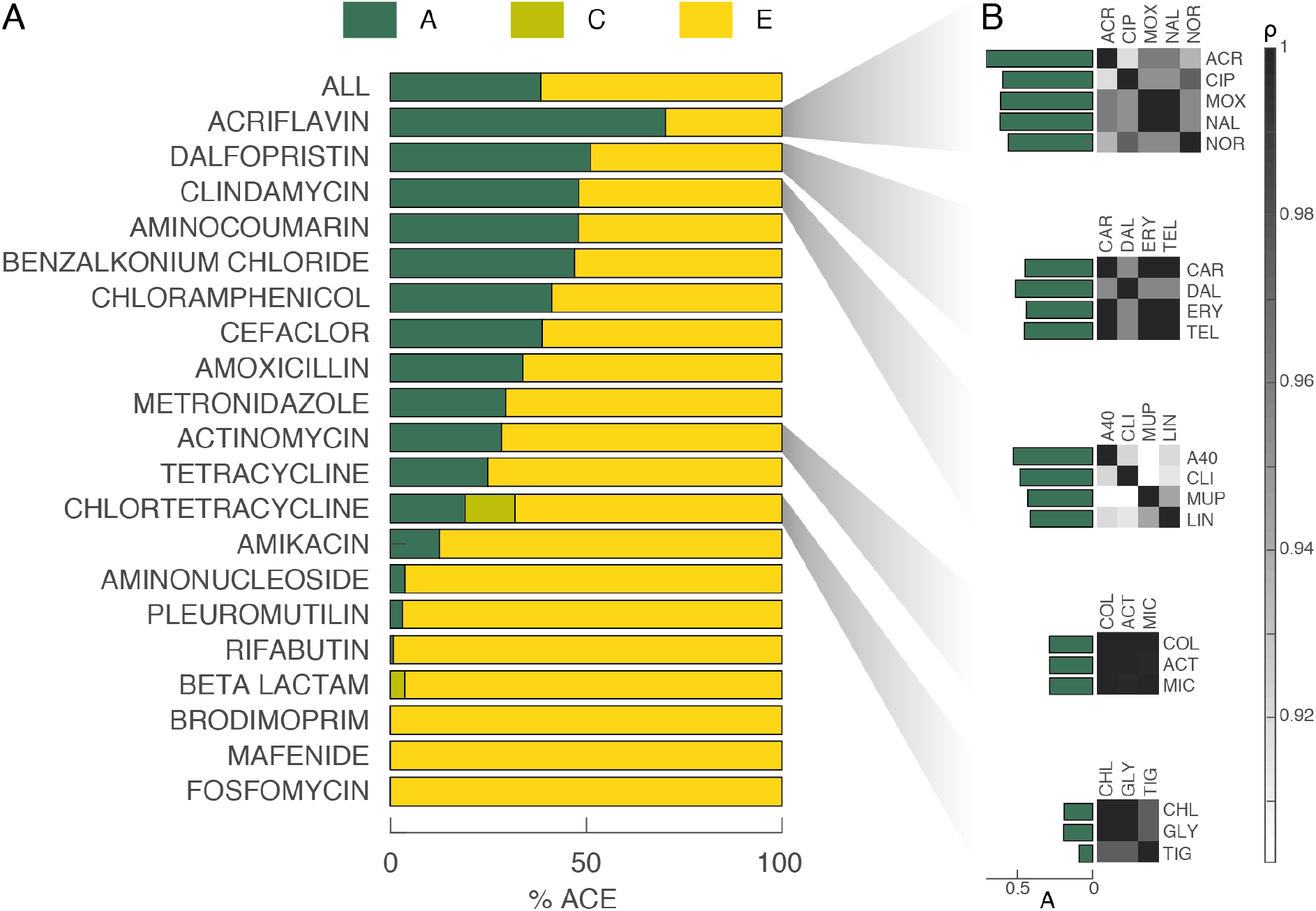
Heritability of the human gut ARP. Heritability estimate results calculated with the OpenMx ACE model. Full results are presented in **Supplementary table 2**. ACR, acriflavin; CIP, ciprofloxacin; MOX, moxifloxacin; NAL, nalidixic acid; NOR, norfloxacin; CAR, carbomycin; DAL, dalfopristin; ERY, erythromycin; TEL, telithromycin; A40, antibiotic a40926; CLI, clindamycin; MUP, mupirocin; LIN, linezolid; COL, colistin; ACT, actinomycin; MIC, microcin J25; CHL, chlortetracycline; GLY, glycylcycline; TIG, tigecycline.

The twin model also allows the decomposition of the environmental variance into components that can be attributed to each individual (E, or unique), or that are shared within a twin pair (C, or common). In our data, the majority of the environmental impacts were attributed to individual-specific effects, in line with previous observations from 16S results [21].

### Heritability of the gut resistome is only partially attributed to taxonomical heritability

Previous studies conducted in the same cohort have showed that the relative abundance of certain taxa in the gut can be heritable [21, 22]. Despite correcting for genus abundance in the ARP calculation, as well as correcting for overall α-diversity in the heritability analyses, it is plausible that the observed genetic contributions to ARPs may be attributed to heritability of different components of the gut microbial community that we may not have corrected for in full. To tackle this, we carried out additional analyses with further corrections specifically for the ARPs that displayed significant proportion of variance explained by host genetics (P < 0.05), namely: acriflavin, aminocoumarin, dalfopristin and clindamycin; as well as the sum of total ARPs (ALL).

Some bacterial genera carry more ARGs on average per genome and will therefore make a greater contribution to an ARP profile. We first evaluated if heritable ARGs were carried by a large number of heritable genera (A > 20%). Xie et al. (2016) reported that in total 27 genera displayed at least moderate heritability (A > 20%) using the same dataset [23]. All 27 genera contribute to total ARP (ALL) and aminocoumarin ARP, while only 19, 8, and 7 of these contributed to clindamycin, dalfopristin and acriflavin ARPs, respectively (**Figure 3A, Supplementary table 4**). In contrast, brodimopim, for which we estimated no heritable components (A=0), showed contribution from only 2 of the 27 these moderately heritable genera.

**Figure 3:**
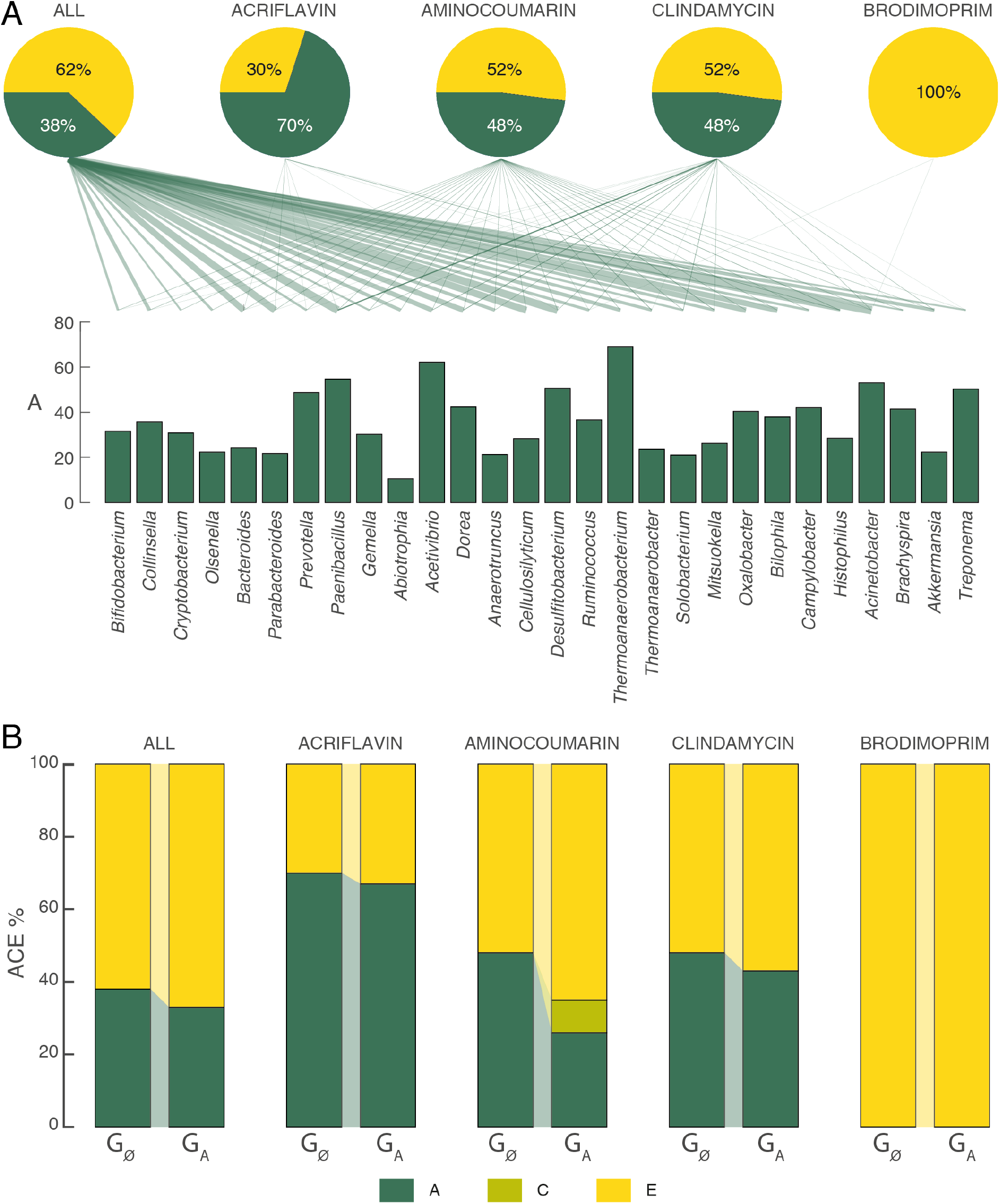
Impact of heritable taxonomy components on ARPs heritability. (A) Link between heritable taxa and the three most heritable ARPs, total ARP (All) and a non-heritable ARP (brodimoprim). Dark green bars represent A (proportion of variance of the trait under genetic influence) estimates previously published for each of 27 heritable genera. The five ARPs represented on the top by pie charts representing their heritability results are linked to genera with A > 0.2 at the bottom. The weight of the link is proportional to the contribution weight of a genus to an ARP. (B) Heritability estimate results for total ARP (All), acriflavin, dalfopristin, aminocoumarin, clindamycin and brodimoprim before (G_Ø_) and after correction for high A (G_A_) bacterial genera relative abundance.

To assess the impact of these genus-level observations on our ARP heritability results, we regressed the four heritable ARPs as well as the sum of all ARPs and brodimoprim (as a negative control) against their contributing heritable genera (A>20%) and used the residuals to re-estimate heritability. The heritability estimates of the sum of all ARPs was reduced by 15% as a result of this correction **(Figure 3B, Supplementary table 5**). For the four other ARPs, we observed a direct relationship between the level of heritability reduction post correction and the number of heritable genera that contributed to each ARP. However, although in all cases the heritability estimates were attenuated after this correction, they still remained nominally significant. The heritability estimates of aminocoumarin ARP, the ARP connected to the greatest number of heritable genera (n = 27) dropped from 48% to 26%. On the other hand, the clindamycin (19 genera), dalfopristin (8 genera), and acriflavin (7 genera) ARPs heritability levels were reduced by only 11%, 0.01% and 4%, respectively, after correction. As expected, the brodimoprim heritability estimate was unaffected by the adjustment.

### The gut resistome is poorly associated with host health status

We next explored if gut resistome profiles are linked to health status of the host in our predominantly healthy older female twin sample. We focused on 24 health traits altogether, including 21 conditions that were reported in at least 10 of the 250 twins, as well as number of days spent in a hospital, visceral fat mass (VFM) estimates, frailty and neuroticism scores (**Supplementary table 1**) and explored their associations with the 21 ARPs using linear mixed effect model adjusted for BMI, alpha diversity, age, gender and family structure. None of the tested associations surpassed FDR at 5% multiple testing correction overall, but allergy and high cholesterol were positively associated with 3 ARPs at FDR 5% correction within health trait (**Figure 4**). Overall, 24 nominally significant associations were observed between 17 health traits and 18 ARPs. These included positive associations between allergy and constipation with 4 and 3 ARPs, respectively, as well as 3 negative associations between VFM and ARPs. In total, 70% of the associations were observed with heritable ARPs. Only three traits (VFM, osteoarthritis and high cholesterol) were associated exclusively with non-heritable ARPs, while eight (frailty, days spent at hospital, UTI, diabetes, thyroid disorders, depression, neuroticism and migraine) were associated exclusively with heritable ARPs.

**Figure 4:**
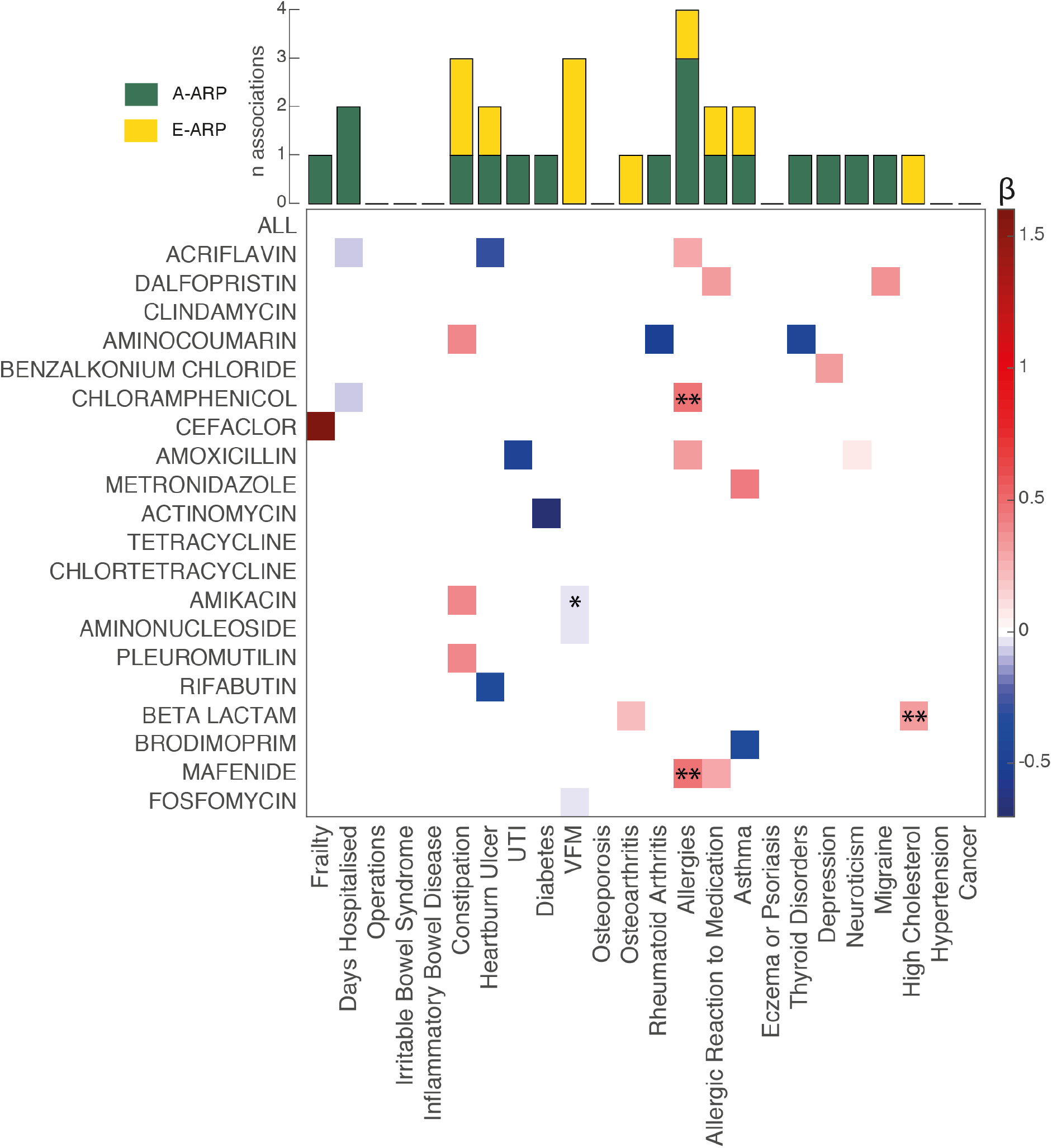
Association between ARP profiles ranked based on their level of heritability and disease. Nominally significant (P < 0.05) associations are colour-coded with blue colours representing negative associations and red colours representing positive ones. * P < 0.01; ** P < FDR 5% for the health condition of interest. The bar graph on the top of the heatmap represents the number of associations observed for each trait with heritable ARPs (A > 20%) in green and with non-heritable ARPs (A < 20%) in yellow. Full results are available in **Supplementary table 6**.

### Host environmental factors are associated with the gut resistome

To evaluate host environmental effects on the gut resistome, we explored the association between ARP variability and 31 host environmental variables, using the linear models as described above (**Supplementary table 1**). Host environment factors were divided into three categories: (i) distal environment, such as the index of multiple deprivation (IMD), has ever lived abroad and contact with pets, (ii) diet, including alcohol consumption and 12 domains calculated from FFQs to build the healthy eating index (HEI), and (iii) medication used by at least 10 participants in the study. Two associations surpassed multiple testing adjustment in the diet category (adjusted P-value < 0.05) and a total of 66 nominally significant associations were observed between 30 environmental factors and 21 ARPs (**Figure 5**).

**Figure 5:**
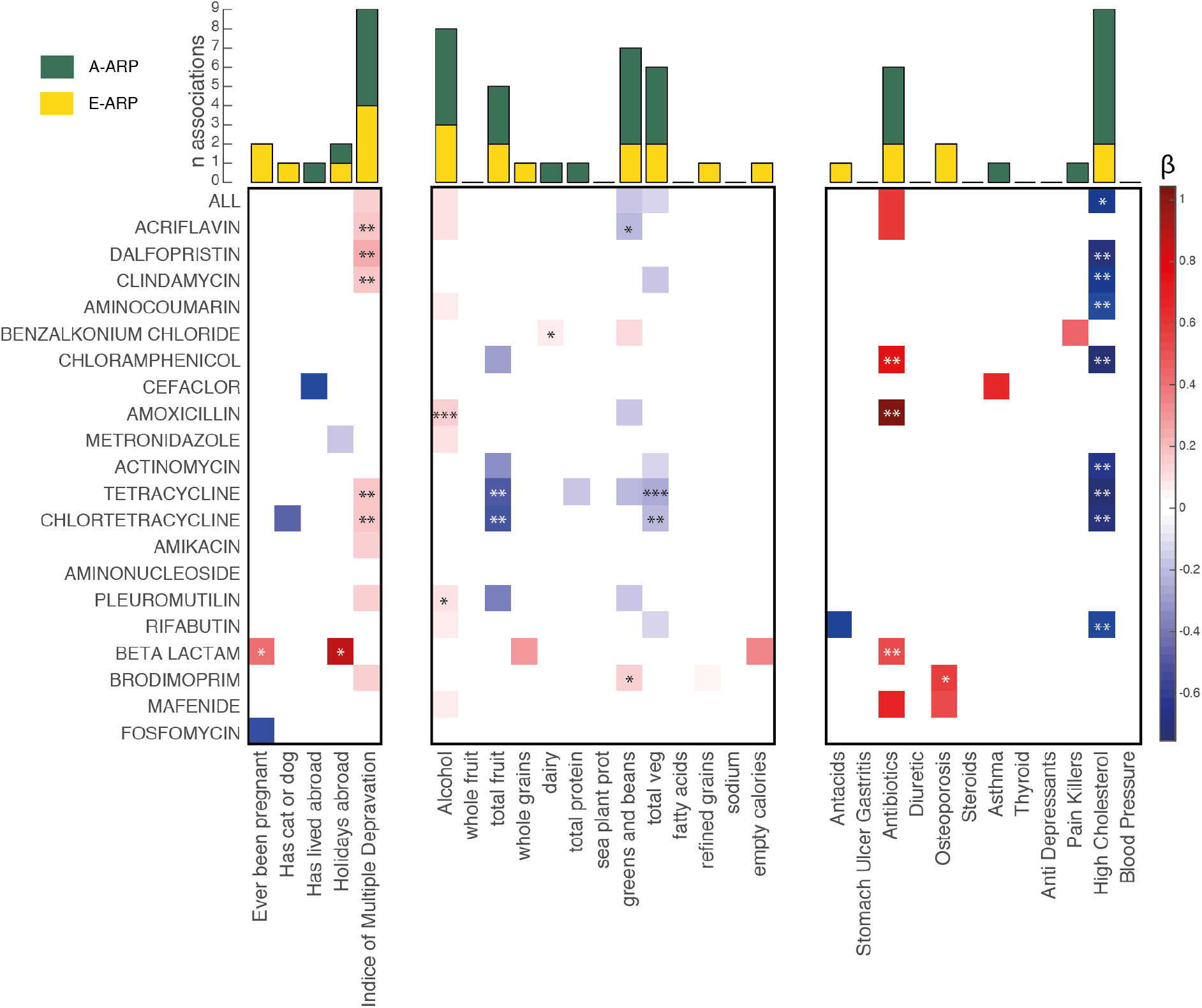
Host environmental factors associate with gut ARP profiles. Association between ARP profiles ranked based on their level of heritability and environmental factors. Nominally significant (P < 0.05) association results are colour-coded on the heatmap with blue colours representing negative associations and red colours positive ones. The bar graph on the top of the heatmap represents the number of associations observed for each trait with heritable ARPs (A > 20%) in green and with non-heritable ARPs (A < 20%) in yellow. * P < 0.01; **P < FDR 5% for individual environmental factors, *** P < FDR 5% within category (distal environment, diet and drugs). Full results are available in **Supplementary table 7**.

Over half of the nominally significant associations were observed with diet variables (31 associations, 54% of total). The most significant results that surpassed multiple testing adjustment were obtained between alcohol consumption and amoxicillin ARP (beta = 0.1561 ± 0.0390; P = 9.58×10^−5^), followed by an association between total vegetables consumption and tetracycline ARP (beta = −0.3840 ± 0.0987; P = 1.44×10^−4^). A large number of nominally significant associations were also observed between ARPs and consumption of greens and beans (associated with 7 ARPs, of which 2 displayed association P-values < 0.01), as well as total vegetables intake (associated with 6 ARPs, of which 1 displayed association P-values < 0.002). Dairy consumption displayed one positive association with benzalkonium chloride ARP (beta = 0.0702 ± 0.0249; P = 0.0053), and total protein consumption was negatively associated with tetracycline ARP (beta = −0.1335 ± 0.0633; P = 0.0380). All associations observed between alcohol consumption and ARPs were positive, while most of those observed between fruit, vegetable or greens and beans consumption with ARPs were negative. For distal environmental factors, there were no results after multiple testing correction, but 15 nominally significant associations were observed, the majority of which were positive associations with IMD (n = 9). Notably, three of the four most heritable ARPs were positively associated with IMD (acriflavin: beta = 0.1983 ± 0.0653, P = 0.0032; dalfopristin: beta = 0.2630 ± 0.0715, P = 0.0005; clindamycin: beta = 0.1799 ± 0.0587, P = 0.0032), as well as sum of all ARPs (beta = 0.1509 ± 0.0706, P = 0.0355). Furthermore, we also observed a positive association between beta lactam and previous pregnancy or pregnancies (beta = 0.4613 ± 0.1389; P = 0.001), as well as having lived abroad (beta = 0.9235 ± 0.3081; P = 0.003). Interestingly, this was the only set of variables for which a majority of the associations (53%) were observed with non-heritable ARPs.

In the medication category, there were no results after multiple testing correction, but 20 nominally significant associations were observed between use of 7 medications and 21 ARPs. As expected, antibiotic consumption was positively associated with 6 ARPs including amoxicillin (beta = 1.0441 ± 0.3179; P = 0.001), beta lactam (beta = 0.5785 ± 0.2028; P = 0.005) and chloramphenicol (beta = 0.8327 ± 0.3019; P = 0.006). However, the highest number of associations (9 associations) were detected with high cholesterol medication, and in all cases, these were negative associations. The strongest association was observed between tetracycline ARP and high cholesterol medication (effect size = −0.75±0.22; P = 0.001). As statins are the most commonly used drugs for high cholesterol, we checked if this signal could be attributed to use of statins. Out of 50 volunteers on high cholesterol medication, 29 (58%) reported statin use. We evaluated the association between ARPs and statin use (excluding volunteers who used high cholesterol drugs other than statins; and using the same mixed effect model as previously described) and observed that none of the 9 associations remained nominally significant. However, for 5 out of 9 associations (ALL, chloramphenicol, chlortetracycline, rifabutin and tetracycline) the direction of the associations remained negative. Thus, it is not possible to exclude that fact that statins use may be the underlying cause of these results, and this would need to be confirmed in a larger study. No concordance was observed between the results obtained for medication use and the corresponding associated condition, where these data were available (**Figure 4**).

## Discussion

We describe the gut resistome of a Caucasian predominantly healthy older female sample from the UK and aim to dissect the role of host genetic and environmental factors on shaping the antibiotic resistance reservoir. Most ARPs were prevalent in over 90% of the population sample, which was much higher than previously reported [10, 11]. This difference is likely due to changes and improvement of the databases used to characterise ARGs and indicates that the majority of the population is likely to harbour many ARGs in the gut, with potential implications for risk of developing resistance to antibiotic treatments in case of infection.

Our results confirm that the human gut resistome is mostly shaped by environmental factors. Yet, we observed that on average over a quarter of ARP variance may be under host genetic control. While some ARPs could be considered not heritable, many (57% of ARPs) were over 20% heritable. The most heritable ARP, acriflavin, showed very strong evidence of host genetic impacts with heritability of 70%. Acriflavin is a topical antiseptic and this observation may be driven by the fact that most common human skin diseases are also heritable [36]. ARP heritability was in line with previous analysis of the TwinsUK microbiome demonstrating that the abundance of both bacterial taxa and genes could be heritable [23]. On the other hand, our estimates of ARPs heritability are greater than expectation based on previously reported host genetic contribution to the taxonomic composition of the gut microbiota [21, 22, 23]. By correcting ARPs for the relative abundance of highly heritable genera that contributed ARGs, we observed a proportion of the measured ARP heritability likely reflects the heritability of the bacterial gene carriers. Nonetheless, this did not fully eliminate the role of host genetics onto the ARP itself. Thus, these results suggest that host genetic effects may not only shape the gut bacterial ecosystem and favour the growth of specific bacterial taxa, but could also promote presence or absence of specific gene functions such as antibiotic resistance within the gut.

The twin model results indicated that, as expected, the majority of antibiotic resistance variation in our sample could be attributed to environmental factors unique to an individual. To explore this further, we compared antibiotic resistance profiles to multiple environmental exposures, lifestyle and health factors. Overall, the strongest and most wide-spread associations were observed with dietary intake components (especially alcohol intake and vegetable consumption), medication use (particularly cholesterol lowering drugs such as statins), and socioeconomic status (SES) defined by the IMD. Dietary components (predominantly alcohol, fruits, vegetables and legume consumption) exhibited associations with multiple ARPs. Diet plays an important role in shaping the gut microbiome [37, 38] that is then able to influence the resistome [39], which may explain our observations. For instance, the HEI built using the dietary components considered here has been strongly associated with the composition of the gut microbiota [34]. Consumption of alcoholic beverages such as red wine has also been associated with alteration of the gut microbiota diversity and composition [40] and may contribute to our results of numerous positive associations observed between alcohol consumption and ARPs in this study. Although positive ARP-associations with alcohol intake was observed, ARP-associations observed with vegetables as well as fruits and beans were all negative. These results could reflect the importance of these foods, potentially through their high fibre content, in modulating the composition of the gut microbiome at the taxonomic level [41, 42], thus affecting the gut resistome. The observed effect of diet on ARPs may also contribute to their associations with SES. Indeed, diet intake has been correlated with SES in numerous studies [43, 44] and we observe here that 9 ARPs were positively associated with IMD, of which 7 are also associated with one of diet items studied. Yet, a recent study demonstrated that the associations detected between IMD and the gut microbiome were not all affected by dietary intake [35]. This suggest that other components of SES may contribute to shaping the gut resistome.

Beside the general effect of diet on the gut resistome, the spread of ARGs across the human population could partly be attributed to the use of antibiotics in the food industry described as the ‘farm-to-fork’ hypothesis [45]. Indeed, it was found that the total ARP levels of a human gut within a country is directly proportional to the quantity of antibiotics use in farms [10]. However, in our study, only two nominally significant associations were observed between ARPs and protein intake suggesting that meat consumption may not be the main driver of ARG transfer. We observed one positive association between dairy consumption and benzalkonium chloride (BC) ARP. BC is an agent that can be used as a disinfectant in the dairy industry, leading to the development of BC resistant bacteria [46, 47, 48]. Furthermore, bacteria from farm animals can be transferred to humans *via* fermented dairy products such as cheese [49]. Together, this suggests that dairy consumption may also be relevant in terms of transfer of ARG from animals to human and selective dietary alteration of the gut microbiota composition at a taxonomic level may also play an important role in shaping the gut resistome. Diet could also contribute to the observed heritability of ARPs as food choices were also described as heritable [50].

We assessed the association between ARPs and medication use as many drugs affect the composition of the gut microbiota [51, 52]. As expected, antibiotic consumption was positively associated with 9 ARPs, as well as the sum of all ARPs, but these results did not surpass multiple testing correction. A recent study following the gut microbiome of 12 men post antibiotic treatment described an increasing trend in ARP levels up to 6 months after exposure [53]. Nevertheless, most of the major effects were observed within a 4 to 8 days window suggesting a time-limited effect of antibiotic consumption on the gut resistome, which may explain our modest association results [53]. The strongest associations were observed with amoxicillin and methicillin ARPs, two commonly used antibiotics, for which resistance of human commensals have been reported [54, 55]. Other drugs had no noticeable effects on the ARP profiles apart from negative associations observed with drugs used to treat high cholesterol. The most common cholesterol lowering drugs are statins, that have been reported to affect the composition of the gut microbiota [51]. Statins have been proposed as potential ‘AMR breakers’, molecules described as capable of re-sensitising bacteria resistant to antibiotics [56, 57]. The observed associations between high cholesterol and ARPs were not significant when considering statin use only, but the lower sample size in the statin subset analyses reduced our power. However, 5 of the 9 associations remained negative, in line with a potential effect of statins on the resistome. While the effect of statins on ARPs would need to be confirmed in a larger sample, our data suggest that other high cholesterol drugs may also be at play and should be studied in more depth.

ARPs were relatively weakly correlated to host health status variables, with only few nominally significant associations observed with common diseases and health-related phenotypes. Surprisingly, the number of urinary tract infections (UTIs) or days spent in the hospital within the last year were negatively associated with ARP levels, despite the fact that hospitals are thought to play a small, but significant role in ARG spread [58, 59], and UTIs are generally eradicated by antibiotic treatment. This result may be due to the small sample size in these analyses, with only 20 volunteers reporting at least 3 UTI in their lifetime, and 19 with a hospital visit (of at least one day) within the last year. On the other hand, autoimmune disorders such as allergy and rheumatoid arthritis, were positively associated with ARPs, in line with our expectations. Both diseases have been associated with alteration of the composition of the gut microbiota [30, 60, 61]. While allergy has been associated with increased *Proteobacteria* that carry a high number of ARGs in our dataset [62], rheumatoid arthritis has mostly been linked to an increase in *Prevotella* [63, 64] that contains less ARGs than the average genus. Interestingly, visceral fat mass was generally negatively associated with ARP levels, which is in line with the negative trend described by Forslund et al. (2014) between ARPs and BMI [11].

Although these results can improve our understanding of the many intrinsic and extrinsic factors that shape the human gut resistome, this study has limitations. First, although this is one of the largest studies of its kind so far, the sample size was relatively limited and only suggestive associations were observed that would need to be replicated in larger samples to lead to robust conclusions. Furthermore, causal mechanisms could not be inferred due to the cross-sectional nature of the study. Most of the phenotypes and environmental exposures that we explored were self-reported, including diet. Ideally future work would explore these findings using objective measures of environmental exposures and diet, and clinically validated phenotypes. Finally, ARPs are an *in-silico* measure of potential for antibiotic resistance, and actual resistance of the gut community would need to be further assessed *in vitro* or *in vivo* to fully assess the impact of host genetic and environmental factors on the resistance of the gut community to antibiotic treatment.

## Conclusions

In summary, our results show that based on our UK female population sample, the human gut can be considered as a reservoir for antibiotic resistance genes. We demonstrated that while the gut resistome is mostly shaped by environmental factors, over a quarter of its variance can be mapped to host genetics and this can only partly be explained by the overall heritability of the gut microbiota composition. Although we are still far from being able to conduct genome-wide association studies that will enable us to understand the role of host or bacterial genetic architecture on the human gut resistome, our results imply that, in the future, host genetic variation could be taken into consideration when prescribing antibiotics. Additionally, we observed that the composition of the human gut resistome is strongly linked to a multitude of environmental factors, beyond antibiotic consumption. Indeed, diet was the environmental component associated with the most ARPs, suggesting that food production and composition may play a key role in the global ARG spread in addition to its effects on the taxonomic composition of the gut microbiome. Altogether, our results suggest that, as for many other therapies, antibiotic prescription should be framed in a personalised context to maximise treatment success and help constrain the spread of antibiotic resistance.

## List of abbreviations

A: additive genetic effects
A40: antibiotic a40926
ACR: acriflavin
ACT: actinomycin
AMR: antimicrobial resistance
ARG: antibiotic resistance genes
ARO: antibiotic resistance ontology
ARP: antibiotic resistance potential
BC: benzalkonium chloride
BMI: body mass index
Bp: bade pair
C: common environment effects
CAR: carbomycin
CHL: chlortetracycline
CI: confidence interval
CIP: ciprofloxacin
CLI: clindamycin
COL: colistin
DAL: dalfopristin
DEXA: Dual-energy X-ray absorptiometry
DNA: Deoxyribonucleic acid
DZ: dizygotic
E: unique environment effects
ERY: erythromycin
FDR: false discovery rate
FFQ: food frequency questionnaires
GLY: glycylcycline
HEI: healthy eating index
IMD: indices of multiple depravation
LIN: linezolid
MIC: microcin J25
MOX: moxifloxacin
MUP: mupirocin
MZ: monozygotic
NAL: nalidixic acid
NOR: norfloxacin
RUC: rural urban classification
SES: socioeconomic status
TEL: telithromycin
TIG: tigecycline
UK: United Kingdome
UTI: urinary tract infection
VFM: visceral fat mass.

## Ethics approval and consent to participate

Ethical approval was granted by the National Research Ethics Service London-Westminster, the St Thomas’ Hospital Research Ethics Committee (EC04/015 and 07/H0802/84). Informed consent was obtained from all volunteer participants.

## Consent for publication

Not Applicable

## Availability of data and materials

The metagenomic shotgun sequencing data for the 250 samples after removal of human sequences reported in this paper are available on the European Bioinformatic Institute (EBI) repository under the following accession number ERP010708. All other phenotypical information’s may be available upon request to the department of Twin Research at King’s College London (http://www.twinsuk.ac.uk/data-access/accessmanagement/).

## Authors’ contributions

J.T.B. conceptualised the study. J.T.B. and S.K.F. supervised the analysis. C.I.LR led the analysis. S.K.F, R.C.E.B, T.C.M, J.C.F, R.C, V.R.C., D.M., C.J.S, and T.D.S. contributed data and analysis inputs. C.I.LR, and J.T.B. wrote the manuscript. All authors reviewed and approved the manuscript.

## Acknowledgements

We acknowledge the study participants from the TwinsUK cohort. We acknowledge support provided by the JPI HDHL funded DINAMIC consortium (http://www.jpi-dinamic.wzw.tum.de/, administered by the MRC UK, MR/N030125/1) and JPI HDHL funded DIMENSION consortium (administered by the BBSRC UK, BB/S020845/1). TwinsUK is funded by the Wellcome Trust, Medical Research Council, European Union, the National Institute for Health Research (NIHR)-funded BioResource, Clinical Research Facility and Biomedical Research Centre based at Guy’s and St Thomas’ NHS Foundation Trust in partnership with King’s College London.

## Supplementary material

**Supplementary Figure 1**: ARPs are highly correlated. Results of the spearman correlation between 41 independent ARPs. ARP clusters are highlighted by the dashed boxes. ARPs selected for the analysis conducted in this study are indicated in bold.

**Supplementary table 1**: Summary statistics of the cohorts and variables used in the study.

**Supplementary table 2**: Full results from the ACE heritability analysis. A, proportion of variance explained by host genetics; C, proportion of variance explained by common environment; E, proportion of variance explained by environment unique to an individual; CI_up, upper 95% confidence interval; CI_low, lower 95% confidence interval; P, p-value.

**Supplementary table 3**: Prevalence of the most heritable ARPs in other European population.

**Supplementary table 4**: Weight of taxa contribution to acriflavin, aminocoumarin, clindamycin, daflopristin, brodimoprim and total (ALL) ARPs. ACE estimates obtained in the Xie et al. publication are also presented [23].

**Supplementary table 5:** Effects of taxonomy on the heritability of acriflavin, aminocoumarin, clindamycin, daflopristin, brodimoprim and total (ALL) ARPs.

**Supplementary table 6:** Association results between ARPs and health parameters obtained using a mixed effect linear model where ARPs were a response and BMI, age, gender and alpha diversity were considered as fixed effects and family structure as a random effect.

**Supplementary table 7:** Association results between ARPs and environmental factors obtained using a mixed effect linear model where ARPs were a response and BMI, age, gender and alpha diversity were considered as fixed effects and family structure as a random effect.

