## Supplementary figures and images for "Host genetic and environmental factors shape the human gut resistome"

### Supplementary figure 1

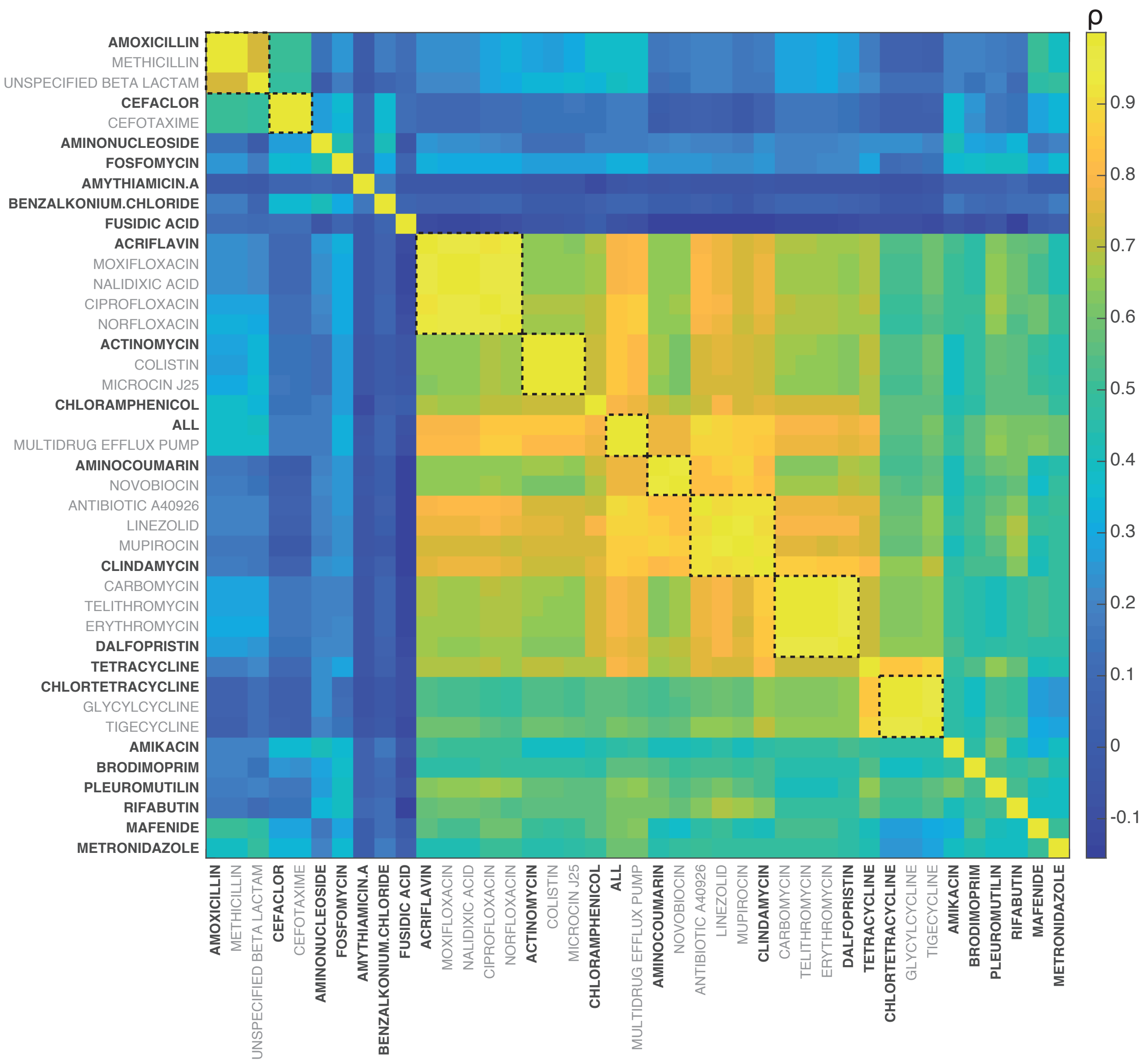
